# FiMO: Inferring the Temporal Order of Mutations on Clonal Phylogeny under Finite-sites Models

**DOI:** 10.1101/2022.01.23.477444

**Authors:** Avesh Kumar Agrawal, Hamim Zafar

## Abstract

Determining the temporal order of somatic mutations that drives cancer progression is essential for understanding the intra-tumor heterogeneity (ITH) and designing personalized therapy. Recently emerged single-cell DNA sequencing (SCS) technologies provide high-resolution datasets suitable for elucidating the temporal order of mutation. However, this task is challenged by technical artifacts associated with single-cell sequencing. While computational methods have been developed to account for these errors, these methods rely on using infinite sites assumption which gets violated in human cancers due to deletion, loss of heterozygosity and parallel mutations. Here, we propose a novel method FiMO, which employs a Bayesian frameowrk for inferring the temporal order of somatic mutations from noisy SCS mutational profiles under tumor evolutionary models that account for mutation recurrence and losses. Using synthetic datasets generated under a wide variety of settings, we show that FiMO outperforms the state-of-the-art methods in inferring the temporal order of mutations. We also applied FiMO on two experimental colon cancer datasets for inferring the temporal order of somatic mutations and quantifying their posterior probability. FiMO is publicly available at https://github.com/aveshag/FiMO.

## 1 Introduction

The progression of tumor is an evolutionary process driven by accumulation and selection of somatic mutations that enable a renegade cell to proliferate more than the surrounding cells [1]. Rapid division of the cell followed by expansion give rise to a tumor clone consisting of identical cells with growth and proliferative advantage which further acquire more somatic mutations and forms genetically diverse sub-clones [2, 3]. The complex interactions between the tumor clones and sub-clones give rise to intra-tumor heterogeneity (ITH) which leads to tumor growth, metastasis, drug resistance, and relapse [4]. Given the important role of the temporal occurrence of mutations in determining the malignancy type [5], it becomes essential to disentangle the temporal order of somatic mutations for elucidating the ITH for developing personalized therapy. Since, genomic data is obtained from tumor at a single snapshot at the time of biopsy, computational methods are required for inferring the temporal order of mutations from DNA sequencing data.

While traditionally bulk sequencing has been the method of choice for profiling somatic mutations from cancer tissue, the mixture of multiple subclones make it difficult to infer temporal mutation occurrence and identify rare subclones [6]. Recently developed single-cell DNA sequencing (SCS) technologies [7] provide high-resolution datasets for comprehensive delineation of ITH and inference of temporal mutation order as genotypes of single tumor cells can be directly measured from single tumor cells. However, the inherent technical artifacts associated with single-cell sequencing protocols including uneven coverage, false negatives (FNs) due to allelic dropout (ADO), and false-positive (FP) errors add additional challenges while determining somatic SNVs and their temporal occurrence [8].

To overcome the challenges due to technical artifacts of SCS while inferring the tumor phylogeny that models the evolutionary process in tumor, several computational methods [9, 10, 11] have been developed. Earlier methods [9] relied on infinite sites assumption that only allows a single occurrence of a somatic mutation. However, cancer evolution is a complex process that harbors events (deletion, loss of heterozygosity (LOH)) that can violate the assumption [12]. To overcome these issues, a finite-sites model of tumor evolution was proposed in the method SiFit [10], which employed a likelihood-based approach for inferring tumor phylogeny while accounting for mutation losses and recurrences. In addition, SiFit can also infer the order of mutations on the branches of the inferred phylogeny. Later, the authors of SiFit further developed a method called SiCloneFit [11] that performs Bayesian inference of clonal phylogeny thus accounting for the cluster structure among the tumor cells. However, the inference of the temporal occurrence of mutations by these methods does not account for the uncertainty in mutation placement on the branches of the phylogeny. Very recently, a method Mutation Order (MO) [13] has been proposed that performs Bayesian inference of temporal order of mutations on a tumor phylogeny derived from SCS data. However, this method works under infinite site assumption and cannot model mutation recurrence and losses.

To overcome these issues, We propose a novel method FiMO (**Fi**nite-sites **M**utation **O**rder), which extends the Bayesian framework of MO for inferring the temporal order of mutations under finite-sites evolution models that account for mutation losses and recurrences. FiMO models the tumor evolutionary process using a finite-sites model that accounts for both parallel and back mutations as well as a Dollo-*d* [14] model that supports for multiple losses of a mutation. Moreover, FiMO also accounts for SCS errors in the Bayesian framework to quantify the uncertainty in the temporal order of mutations along the branches of a clonal phylogeny leaves of which represent tumor clones. The framework of FiMO is general and also supports cell-level phylogeny. Using synthetic datasets under varied experimental settings, we show that FiMO outeprforms MO and SiFit’s mutation order inference based on multiple metrics. In order to facilitate the comparison of temporal mutation order where the inferred phylogeny may differ from the ground truth, we also introduce a new metric called partition accuracy. Finally, we applied FiMO on two experimental datasets from colon cancer patients to derive the temporal order of mutations and quantify the posterior probability of mutation placement on the clonal phylogeny derived from SiCloneFit.

## 2 Methods

Given a tumor phylogeny (with branch lengths) that represents the genealogical relationship between *m* single cells sampled from the tumor, the goal of FMO is to determine the branches where the somatic mutations occurred on the tumor phylogeny assuming a finite-sites tumor evolution model. We further assume that evolution of each somatic mutation site is independent and the SCS technical errors independently affect each site. The input to FMO consists of an *n* × *m* binary genotype matrix denoting the observed genotypes of *m* single cells for *n* somatic mutation sites, a clonal phylogeny 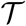 with *k* leaves representing *k* clones and the membership of cells to clones, false positive (FP), false negative (FN) and recurrent mutation rates and the maximum number of times a mutation can be lost (*d*) for Dollo-*d* model.

### 2.1 Notation and Terminology

In this work, we consider binary SCS mutation data where m single cells have been sampled from different tumor clones and for a mutation site, 0 and 1 denote the absence and presence of the mutation respectively. Each of *n* mutation sites evolve along the clonal phylogeny 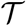 whose leaves represent tumor clones *CN*_*i*_, where *i* = {1, 2, …*k*}.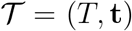 consists of a tree topology *T* and a branch length vector **t**. The tree topology *T* = (*V*, *E*) is a rooted binary tree with *V* denoting the set of nodes and *E* = {*e*_1_, *e*_2_, …e_2*k*-2_} denoting the set of branches ((2k − 2) branches for a binary tree with *k* leaves). The root *r* of *T* denotes a normal cell lacking any somatic mutation.

Let *e* ∈ *E* is a branch that connects two nodes *v* and *w* with *v* as the immediate ancestor. We define *U*^*b*^(*w*) as the clade induced by *w* and contains the node *w* and all its descendent nodes. Branch *b* denotes the *ancestor branch* of clade *U*^*b*^(*w*). The set of branches (subset of *E*) within the clade *U*^*b*^(*w*) is denoted by *E*^*b*^(*w*), and *L*^*b*^(*w*) denotes the set of leaves in clade *U*^*b*^(*w*). Let *ancs*(*b*) denote the set of ancestor branches of *b*. Let 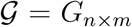 and 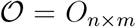 denote the true genotype matrix and observed genotype matrix for *n* somatic mutation sites and *m* single cells respectively. *O*_*ij*_, the (*i*, *j*)^*th*^ entry in 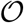 can differ from *G*_*ij*_, the (*i*, *j*)^*th*^ entry in 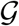 because of SCS technical errors.

### 2.2 Somatic Mutation Process

To model the somatic mutation process in cancer for binary data, we adopt two different models - 1) a finite-sites model adopted from SiFit [10] that accounts for the effects of mutation losses and recurrence, and 2) a Dollo model that accounts for mutation losses.

#### 2.2.1 Finite-sites Model

The finite-sites model of SiFit [10] uses a continuous-time Markov chain for modeling possible transitions of the genotypes. Using this, we model the mutation process for branch length *t* by a 2 × 2 transition probability matrix *P*(*t*) which is obtained by matrix exponentiation of the product of transition rate matrix *Q* of the Markov chain and the branch length *t*. The product of the transition-rate matrix and the branch length *t* is given by

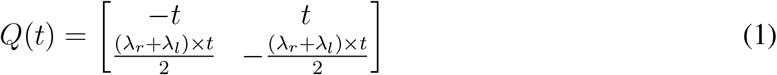

where λ_*r*_ and λ_*l*_ accounts for the effects of recurrent mutation and mutation loss respectively.

The finite-sites model include three possibilities for a mutation *i* occurring on a branch *b* in 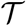:

1. Single Mutation: A branch *b* harbors the mutation (0 → 1) and no other branch is mutated for this site, represented by the pair (*b*, *Φ*), where *Φ* is a null branch
2. Back Mutation: The mutation first occurs (0 → 1) on a branch *b* and then one of its descendent branch (*y* ∈ *E*^*b*^(*w*)) harbors a loss of mutation (1 → 0), denoted by the pair (*b*, *y*)
3. Parallel Mutation: A site *i* is mutated at two different branches (*b* and *y*) in two different sub-trees (0 → 1 and 0 → 1), involves a pair of branches (*b*, *y*)

Assuming mutation *i* to occur on branch *B*_*i*_ = *b*, 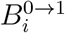 denotes a branch where mutation site *i* is mutated from 0 to 1 and 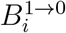 denotes a branch (∈ *E*^*b*^(*w*)) harboring back mutation. The marginal probabilities of occurrence of mutation *i* on a branch *b* for all three possible conditions are given by

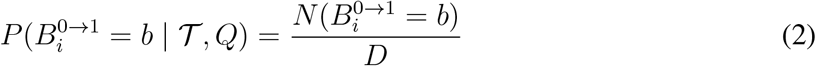

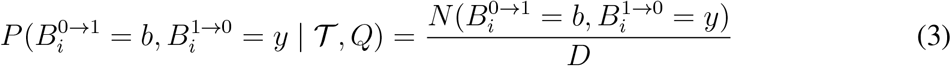

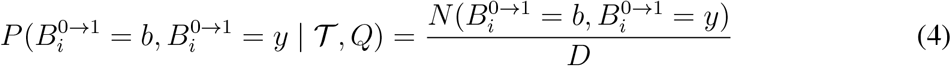

Where

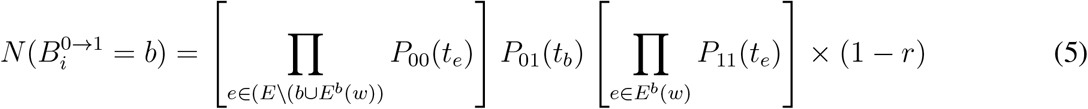

In eq. (5), *t*_*e*_ is the length of branch *e* and *r* is the prior probability of a recurrent mutation. The first term is a product of probabilities across all branches where mutation did not occur, the second term is the probability of mutation occurring on branch *b*, and the third term is a product of probabilities across all branches in *E*^*b*^(*w*).

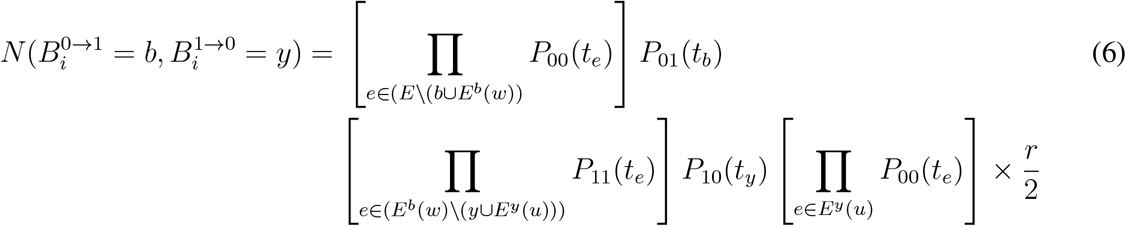

In eq. (6), the first term is a product of probabilities across all branches where mutation did not occur, the second term is the probability of mutation occurring on branch *b*, the third term is a product of probabilities across all branches in *E*^*b*^(*w*) prior to the back mutation occurring on branch *y*, the fourth term is the probability of the back mutation occurring on branch *y*, and the fifth term is a product of probabilities across all branches in *E*^*y*^(*u*) (descendants of back mutated branch) without the mutation.

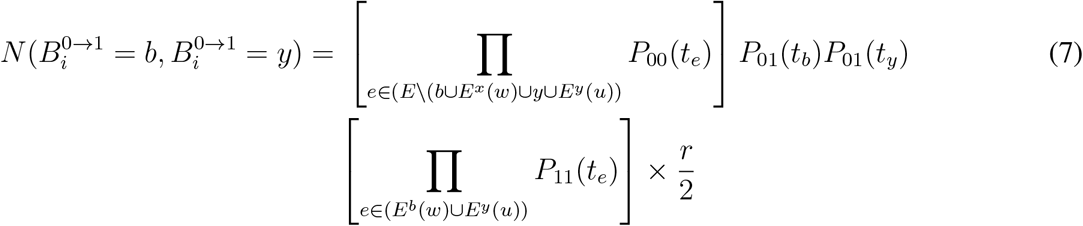

In eq. (7), the first term is a product of probabilities across all branches where mutation did not occur, the second and third terms are the probabilities of the mutation occurring on branches *b* and *y*, i.e., parallel mutation, the fourth term is a product of probabilities across all branches in *E*^*b*^(*w*) and *E*^*y*^(*u*).

The denominator *D* is required to create a valid probability distribution across all possible pairs of branches, and is calculated by summing all possible numerators across all valid pairs (*z*_1_, *z*_2_) of branches *z*_1_ ∈ *E* and *z*_2_ ∈ *E* ∪ *Φ*:

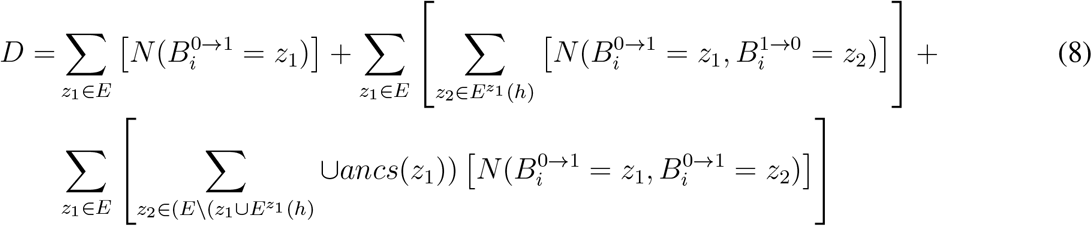

An example of the marginal probability calculation for single, back and parallel mutations under the finite-sites model is given in Supplementary Methods.

#### 2.2.2 Dollo Model

Next we considered the Dollo-*d* model where a mutation can occur exactly once but can be lost at most d times [14]. The Dollo model for binary data include two possibilities for a mutation *i* occurring on a branch *b* in 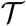:

1. Single Mutation: A branch *b* harbors the mutation (0 → 1) and no other branch is mutated for this site, represented by the pair (*b*, *Φ*), where *Φ* is a null branch
2. Back Mutation: The mutation first occurs (0 → 1) on a branch *b* and then at most d of its descendent branches 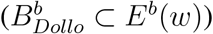 harbors a loss of the mutation (1 → 0)

Let *B*_*i*_ = *b* is a branch where mutation *i* occurs then 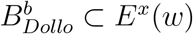 denotes a set of branches where back mutation occurs for mutation *i* under the Dollo model. 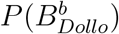 denotes a set of all different possible sets where back mutation can occur and it does not contain null set. The marginal probability that mutation *i* occurs on a branch *b* for above two possible scenarios is given by

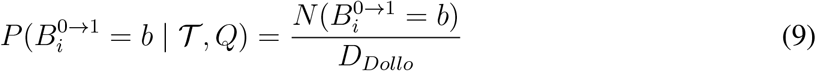

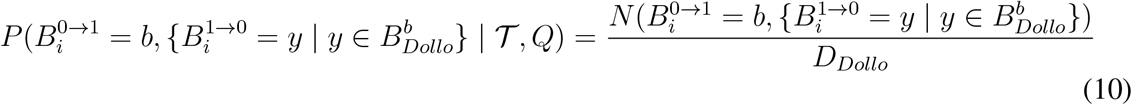

Where 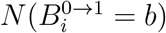 is obtained from eq. (5) and the numerator in eq. (10) is given by

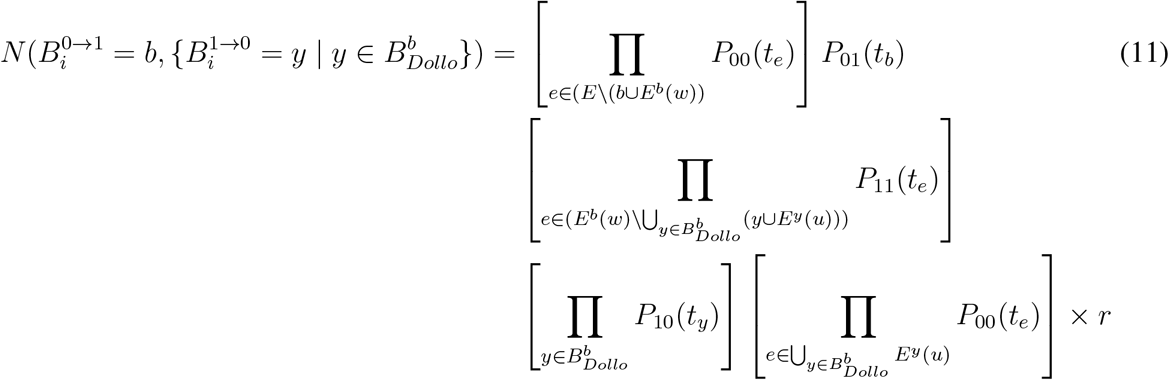

In eq. (11), the first term is a product of probabilities across all branches where mutation did not occur, the second term is the probability of mutation occurring on branch *b*, the third term is a product of probabilities across all branches in *E*^*b*^(*w*) excluding all the branches where back mutation occurred, the fourth term is a product of probabilities across all branches where mutation loss (back mutation) occurred 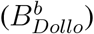, and the fifth term is a product of probabilities across all branches without the mutation that are descendants of back mutated branches (all branches in 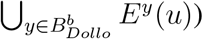. The denominator *D*_*Dollo*_ is given by

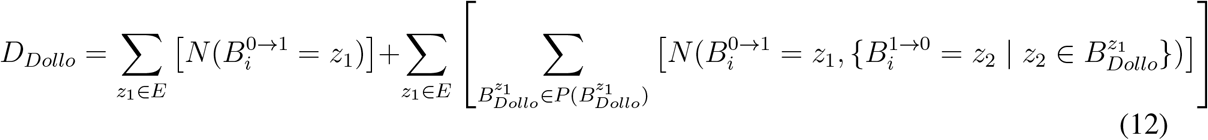

An example of the marginal probability calculation under the Dollo model is given in Supplementary Methods.

### 2.3 Quantification of SCS errors

To quantify the effects of errors in SCS data, we adopt the error model for binary data from the existing works [9, 10]. Assuming *α*_*ij*_ be the false positive rate and *β*_*ij*_ be the false negative rate for genomic site *i* for cell *c*_*j*_, the conditional probabilities of the observed genotype given the true genotype at genomic location *i* of cell *c*_*j*_ are given by:

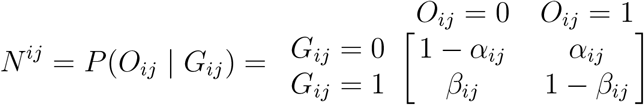

where 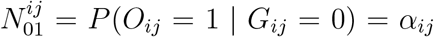 and so on. Given the independence of sequencing errors, if mutation *i* occurs on branch *b*, we can quantify the impact of SCS technical errors for mutation *i* as

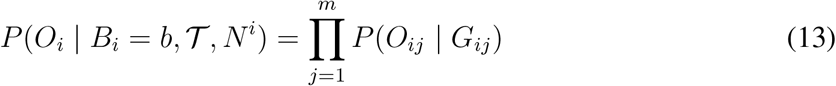

An example quantification of SCS error is given in the Supplementary Methods.

### 2.4 Inferring the location of a mutation in 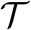

To calculate the posterior probability distribution 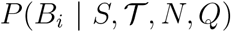 for the location of mutation *i* by using Bayes’ Theorem, we use the following equation

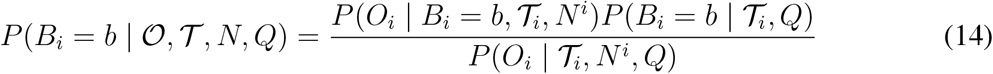

In eq. (14), 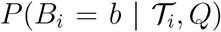 can be calculated by summing over all the scenarios that involves the occurrence of mutation *i* on branch *b*. For the finite-sites model, it is calculated by

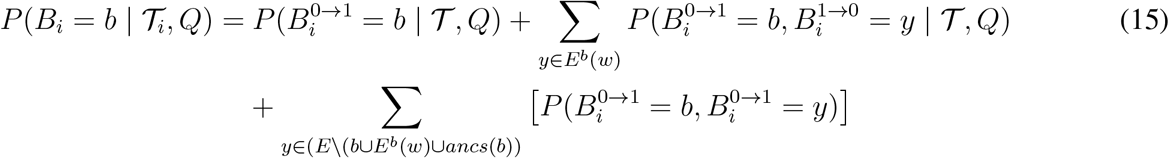

For the Dollo-*d* model, it is calculated by:

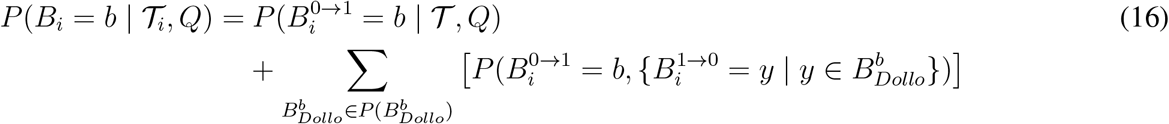

Branch that maximizes the posterior probability is selected as the location for a mutation, i.e., the maximum a posteriori (MAP) estimate. The MAP calculation for the location of mutation *i* is given by

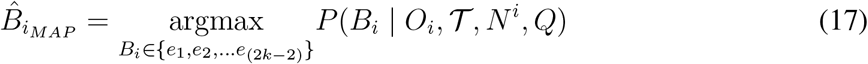

## 3 Results and Discussion

### 3.1 Simulation Study

To evaluate the performance of our method FiMO in predicting the mutation locations, we generated synthetic datasets under different settings: 1) varying the number of cells, 2) varying the FN error rate, 3) varying the value of *d* for the Dollo Model (see Table S1 for details), and 4) varying the number of clones. The simulation process is described in Supplementary Methods and involves the finite-sites evolution model for generating true clonal genotypes. For each experimental setting, 5 replicates were simulated. The accuracy of FiMO and the competing methods (MO and SiFit’s mutation placement algorithm) for each setting was measured using four different metrics. We adopted the metrics location accuracy, order accuracy, adjacent order accuracy from [13]. In addition, we designed a new metric called partition accuracy which is designed for the scenario when the inferred phylogeny differs from the true phylogeny. Partition accuracy measures the accuracy of mutated and non-mutated cells after the placement of a mutation on the inferred phylogeny. All metrics are described in the Supplementary Methods.

In the first scenario, we investigated the performance of FiMO in comparison to other methods when the number of cells was varied (*m* = 100 and *m* = 500). Number of mutation sites was set to 50, number of clones was set to 15 and FN rate = 0.2 was used. Under each experimental setting, FiMO performed the best in terms of all four metrics fig. 1. For both FiMO and SiFit’s mutation placement algorithm, each accuracy measure improved with an increase in number of cells. In contrast, for MO, only adjacent order accuracy improved for the datasets with larger number of cells, other accuracy measures remained similar. For 500 cell datasets, FiMO achieved near perfect accuracy for all four metrics. Next, we evaluated how FiMO performs when the error rates increase. For that we simulated 100 and 500 cells datasets while FN error rate was set to 0.2 and 0.3. Accuracy of each method decreased with increase in error rate under each setting. However, FiMO outperformed the other methods based on all four accuracy metrics under each experimental setting (Supplementary Results, Figs. S10-S11).

**Figure 1:**
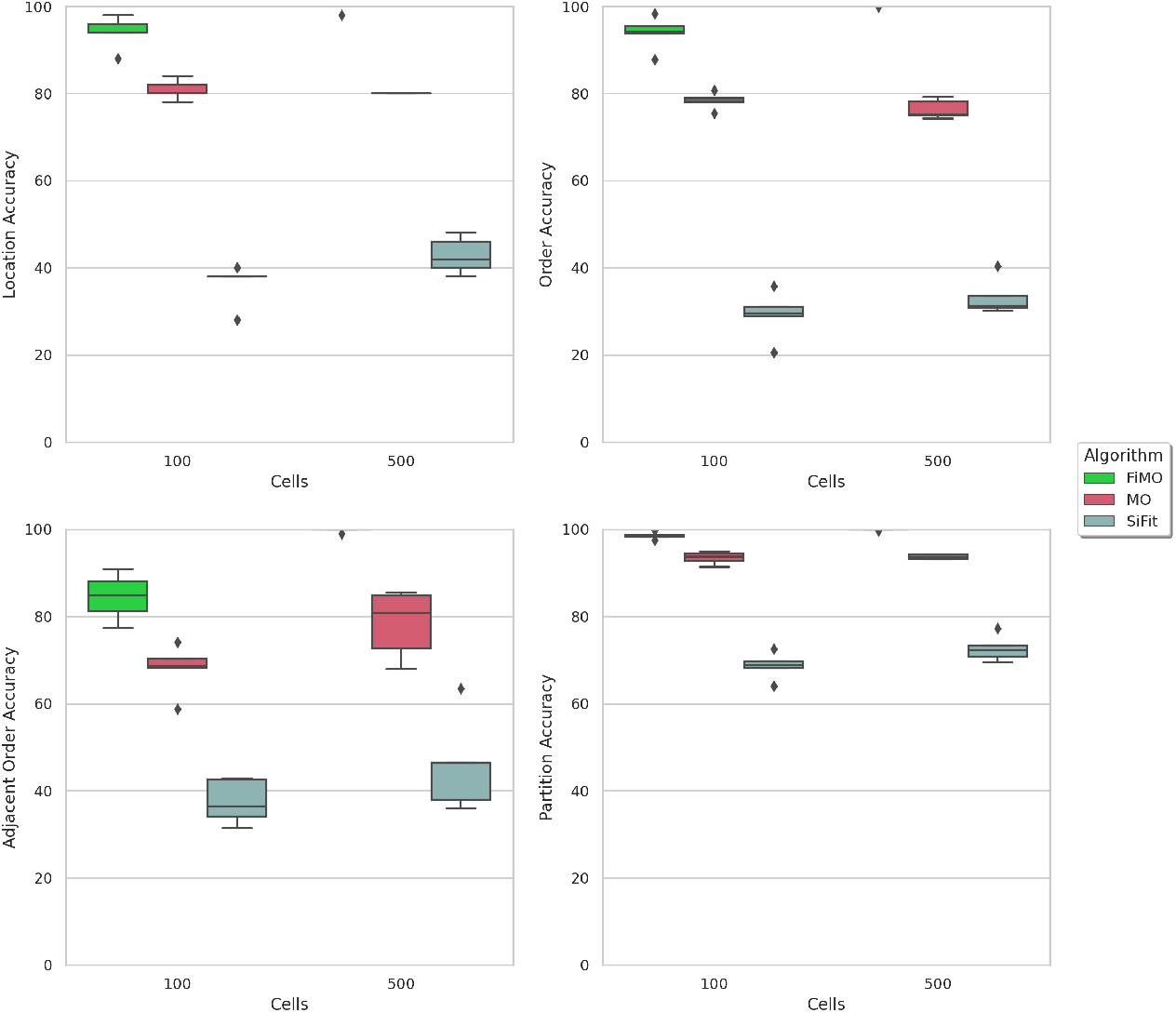
Performance comparison of FiMO, MO and SiFit for varying number of cells. Number of cells was varied *m* ∈ {100, 500}. The y-axes show different accuracy measures.

Next, we evaluated the performance of FiMO while using the Dollo model by generating 100 cells datasets and varying the value of *d* ∈ {2, 3}. With an increase in *d*, performance of each method drops as can be seen from all metrics (Fig. S12). Again under each setting, FiMO performed the best. For datasets with varying number of clones, FiMO performed the best under each scenario (Fig. S13). We also generated datasets under infinite sites assumption for which FiMO and MO performed identically and outperformed SiFit (Fig. S14). Average accuracy of FiMO for each setting is given in Table S2 of Supplementary Results.

### 3.2 Application to Real Datasets

To evaluate FiMO on experimental SCS datasets, we used two colon cancer single-cell DNA sequencing datasets where primary as well as metastatic tumor cells were sampled for both patients [15]. Each of these datasets contain a large number of cells and less number of mutation sites as targeted sequencing was performed.

To prepare the inputs for FiMO, we applied SiCloneFit [11] for inferring the clonal phylogeny. The obtained clonal tree was rooted by adding the root on the branch that separated the normal cell clone from the other tumor clones harboring somatic mutations. SiCloneFit also inferred the FP and FN rate and recurrent and back mutation probabilities. All these parameters along with the inferred phylogeny and the observed genotype matrix were used as input for FiMO.

The first dataset consisted of 178 cells for which 16 somatic SNVs were profiled. SiCloneFit inferred a clonal phylogeny with five clones - two subclones of primary tumor cells, one subclone of metastatic aneuploid tumor cells, a subclone of diploid metastatic cells and a cluster of normal cells that lacked any somatic mutations (fig. 2). This clonal tree along with the FP rate=0.012, FN rate=0.136, recurrent and back mutation probability = 0.1 as estimated by SiCloneFit were used for executing FiMO on this dataset.

**Figure 2:**
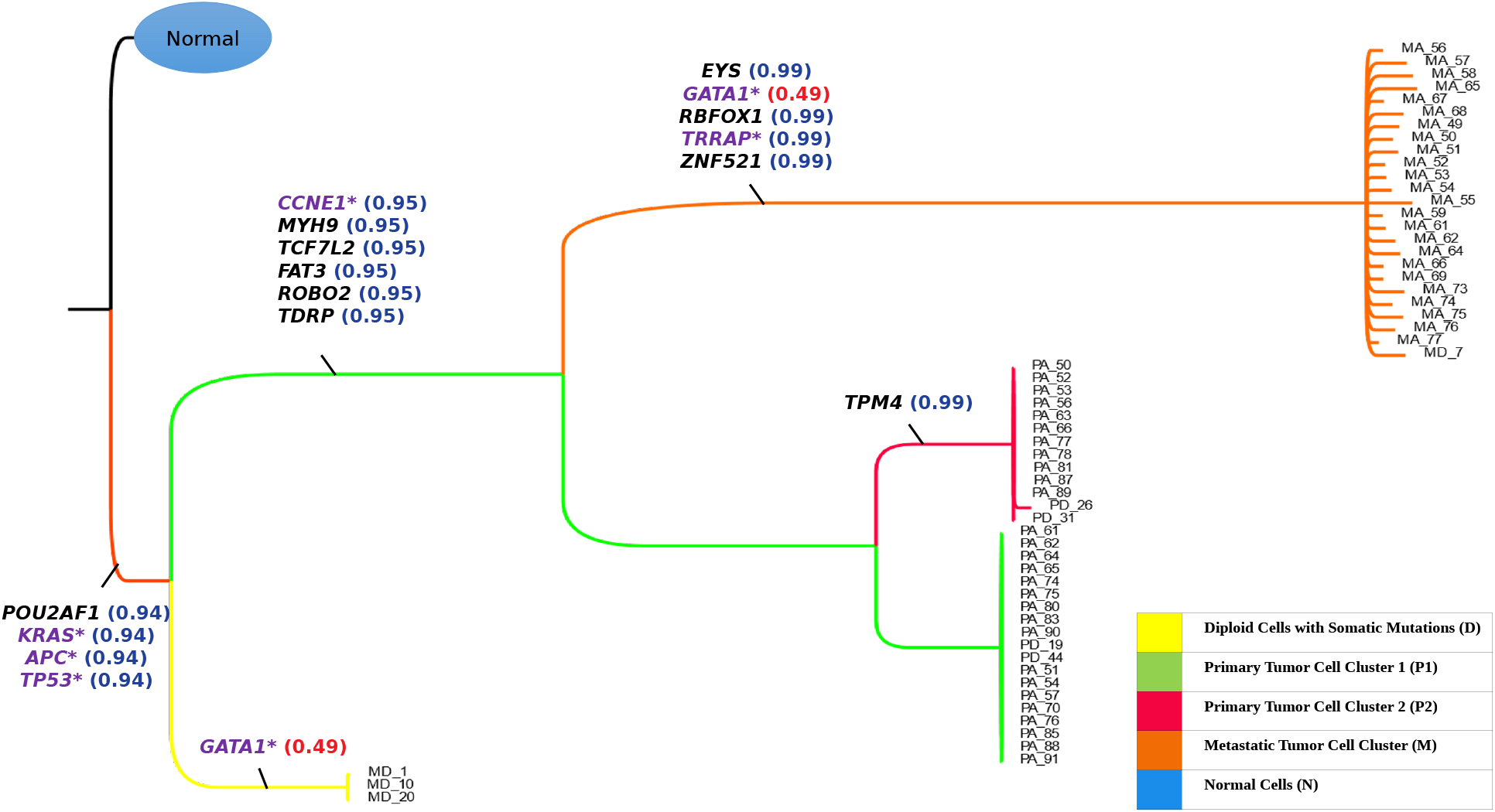
Inferred clonal phylogeny and the temporal order of mutations inferred by FiMO for a metastatic colorectal cancer patient CRC1. SiCloneFit was used to infer tumor clones and clonal phylogeny. Each branch is annotated by somatic mutations whose occurrences are inferred by FiMO. The purple mutations are cancer genes and tumor-suppressor genes. Also shown are the posterior probabilities of mutation locations as inferred by FiMO.

The mutation locations as inferred by FiMO are shown in fig. 2. The formation of the tumor was driven by mutations in *APC*, *KRAS* oncogene and the *TP53* tumor suppressor gene. The primary tumor subclones formed after acquiring six additional somatic mutations that included a mutation in the *CCNE1* oncogene. The metastatic clone evolved from this branch by acquiring mutations in 5 genes including *GATA1* and *TRRAP*. The primary subclone evolved further and acquired a mutation in *TPM4*. The diploid metastatic cells diverged before the occurrence of the mutations in primary tumor cells and was driven by a parallel mutation in *GATA1*. All the mutation locations were supported by high posterior probability. Since *GATA1* had a parallel mutation, the posterior probability was 0.49 on two branches. Thus FiMO was able to quantify the posterior supports for the mutation occurrences.

The second data set had 182 cells that harbored 36 somatic SNVs. SiCloneFit inferred a clonal phylogeny with 6 clones - a normal cell cluster devoid of somatc mutations, two clusters of primary aneuploid tumor cells (P1 and P2), two clusters of metastatic aneuploid tumor cells (M1 and M2), and a cluster (I) harboring diploid cells that contained somatic mutations different from the primary or metastatic clusters. This clonal phylogeny along with the FP rate = 0.012, FN rate = 0.295, mutation recurrence and loss probability = 0.1 as estimated by SiCloneFit were used for inferring mutation locations using FiMO.

The mutation locations inferred by FiMO are shown in fig. 3. *TOX* was a clonal mutation present in all the cells containing somatic mutations. After that the primary tumor clone P1 and the independent clone I diverged. The primary tumor clone P1 acquired mutations in *APC, NRAS, CDK4,* and *TP53* along with 4 other somatic mutations. On the other hand, the clone I (7 diploid cells) acquired mutations in *SPEN, ALK, ATR, NR3C2,* and *EPHB6* that were not shared with any other tumor cells. After acquiring another mutation in *APC* along with two other mutations, the primary tumor gave rise to first metastatic subclone (M1) which acquired 10 metastasis-specific SNVs. Two more mutations (one in *FHIT*) were acquired before the divergence of the second metastatic subclone (M2) and these two mutations were shared by M2 and second primary tumor subclone P2. P2 kept acquiring mutations in *LRP1B* and *LINGO2* that did not occur in any of the metastatic clones. M2 acquired 7 more metastatic-specific mutations, 3 of which were parallel occurrence of mutations in *PTPRD, FUS,* and *LINGO2* that also occurred in M1. All the mutation locations were supported by high posterior probability. The posterior probability for the parallel mutations were divided equally between the two branches.

**Figure 3:**
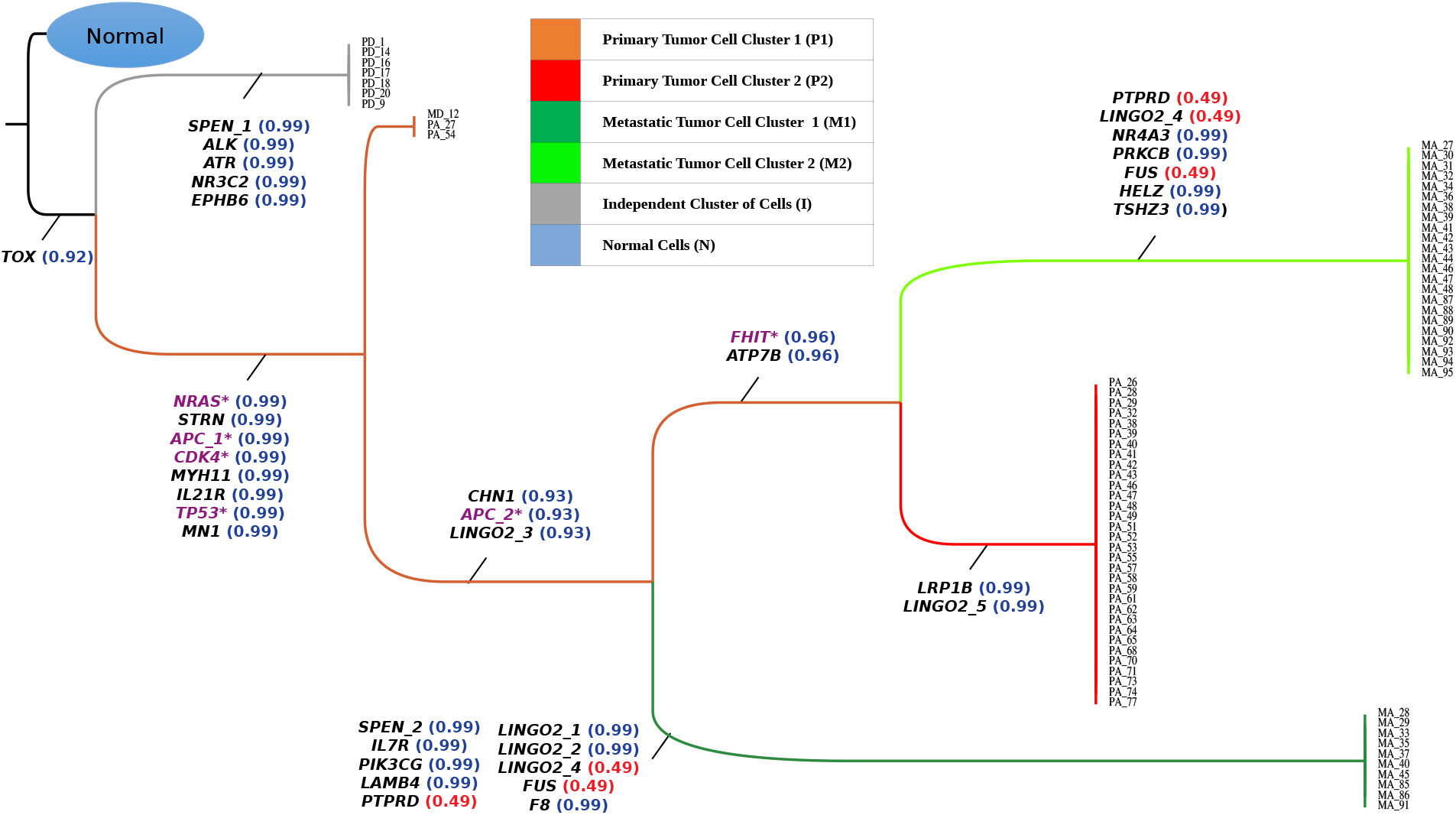
Inferred clonal phylogeny and the temporal order of mutations inferred by FiMO for a metastatic colorectal cancer patient CRC2. SiCloneFit was used to infer tumor clones and clonal phylogeny. Each branch is annotated by somatic mutations whose occurrences are inferred by FiMO. The purple mutations are cancer genes and tumor-suppressor genes. Also shown are the posterior probabilities of mutation locations as inferred by FiMO.

## 4 Conclusion

Temporal order of mutations in cancer progression play a crucial role in providing insight into mechanism that lead to ITH. With the advent of single-cell sequencing techniques, we have high-resolution data to elucidate the temporal order of somatic mutation. However, the errors associated with SCS data necessitates the development of computational methods that can acocunt for errors and uncertainty at multiple levels. Here, we propose FiMO, a novel method that allows the Bayesian inference of temporal order of mutations under finite-sites tumor evolution models that account for the possible losses and recurrences of mutations.

We benchmarked the performance of FiMO using a wide range of synthetic datasets where we varied the number of cells, number of clones, error rates, and tumor evolution model. Under each experimental setting, FiMO outperformed the state-of-the-art methods MO and SiFit. To enable the comparison of temporal order of mutations when the inferred phylogeny differs from the ground truth, we introduced a new metric called partition accuracy. FiMO outeprformed the other methods based on this new metric as well as three other state-of-the-art metrics for evaluating temporal order of mutations along the branches of a phylogeny. By applying FiMO on two experimental datasets from colon cancer, we quantified the posterior probabilities of mutation locations on the branches of clonal phylogenies as inferred by SiCloneFit. FiMO was able to determine the mutation locations with high posterior probability and identify the recurrences of somatic mutations as well as compute their posterior support.

FiMO’s framework is general and can be extended for utilizing the information on missing data to better quantify the uncertainties. FiMO can be improved by integrating models for copy number alterations, as well as by taking into account mutations that affect the same allele multiple times and we plan to pursue these in the future. Although we focused on cancer, FiMO’s framework can easily be extended to work with single cell mutation profiles from a wide range of fields by appropriate modeling of the evolutionary process.

## References

[1] Bert Vogelstein, Nickolas Papadopoulos, Victor E. Velculescu, Shibin Zhou, Luis A. Diaz, and Kenneth W. Kinzler. Cancer genome landscapes. Science, 339(6127):1546–1558, 2013.

[2] PC Nowell. The clonal evolution of tumor cell populations. Science, 194(4260):23–28, 1976.

[3] Lauren M.F. Merlo, John W. Pepper, Brian J. Reid, and Carlo C. Maley. Cancer as an evolutionary and ecological process. Nat Rev Cancer, 6(12):924–935, Dec 2006.

[4] Robert J. Gillies, Daniel Verduzco, and Robert A. Gatenby. Evolutionary dynamics of carcinogenesis and why targeted therapy does not work. Nat Rev Cancer, 12(7):487–493, Jul 2012.

[5] Christina A. Ortmann, David G. Kent, Jyoti Nangalia, Yvonne Beata Silber, David C. Wedge, Jacob Grinfeld, E. Joanna Baxter, Charles E Massie, Elli Papaemmanuil, Suraj Menon, Anna L. Godfrey, Danai Dimitropoulou, Paola Guglielmelli, Beatriz Bellosillo, Carles Besses, Konstanze Döhner, Claire N Harrison, George S. Vassiliou, Alessandro Maria Vannucchi, Peter J. Campbell, and Anthony R Green. Effect of mutation order on myeloproliferative neoplasms. The New England journal of medicine, 2015.

[6] Nicholas Navin. Cancer genomics: one cell at a time. Genome Biology, 15(8):452–465, 2014.

[7] Charles Gawad, Winston Koh, and Stephen R. Quake. Dissecting the clonal origins of childhood acute lymphoblastic leukemia by single-cell genomics. Proceedings of the National Academy of Sciences, 111(50):17947–17952, 2014.

[8] Hamim Zafar, Nicholas Navin, Luay Nakhleh, and Ken Chen. Computational approaches for inferring tumor evolution from single-cell genomic data. Current Opinion in Systems Biology, 7:16–25, 2018. • Future of systems biology • Genomics and epigenomics.

[9] Katharina Jahn, Jack Kuipers, and Niko Beerenwinkel. Tree inference for single-cell data. Genome biology, 2016.

[10] Hamim Zafar, Anthony Tzen, Nicholas Navin, Ken Chen, and Luay Nakhleh. SiFit: inferring tumor trees from single-cell sequencing data under finite-sites models. Genome Biology, 18(1):178, Sep 2017.

[11] H. Zafar, N. Navin, K. Chen, and L. Nakhleh. SiCloneFit: Bayesian inference of population structure, genotype, and phylogeny of tumor clones from single-cell genome sequencing data. Genome Research, 29:1860–1877, 2019.

[12] Alexander Davis and Nicholas E. Navin. Computing tumor trees from single cells. Genome Biology, 17(1):1–4, 2016.

[13] Yuan Gao, Jeffrey Gaither, Julia Chifman, and Laura Salter Kubatko. A phylogenetic approach to inferring the order in which mutations arise during cancer progression. bioRxiv, 2020.

[14] Simone Ciccolella, Camir Ricketts, Mauricio Soto Gomez, Murray Patterson, Dana Silverbush, Paola Bonizzoni, Iman Hajirasouliha, and Gianluca Della Vedova. Inferring cancer progression from Single-Cell Sequencing while allowing mutation losses. Bioinformatics, 2020.

[15] Marco L. Leung, Alexander Davis, Ruli Gao, Anna K. Casasent, Yong Wang, Emi Sei, Eduardo Vilar, Dipen M. Maru, Scott Kopetz, and Nicholas E. Navin. Single-cell dna sequencing reveals a late-dissemination model in metastatic colorectal cancer. Genome research, 2017.

