## Supplementary Material for "FiMO: Inferring the Temporal Order of Mutations on Clonal Phylogeny under Finite-sites Models"

January 14, 2022

#### 1 Supplementary Methods

##### 1.1 An example of Clade

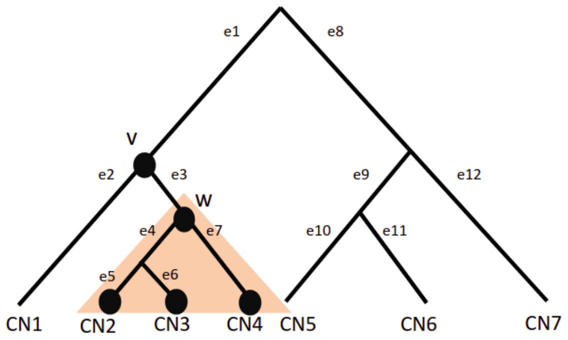


Fig. S1: Shaded triangle is a Clade $U^b(w)$ induced by $w$ with $E^b(w) = \{e4, e5, e6, e7\}$ and $L^b(w) = \{CN2, CN3, CN4\}$

##### 1.2 Example of marginal probability calculation for finite-sites model

An example of the marginal probability calculation for single, back and parallel mutations are given below.

Fig. S4 and fig. S5 showing the observed and true binary genotypes for mutations as illustrated in tree (fig. S2). The set of branches is $E = e_1, ..., e_{12}$ and the corresponding set of branch lengths would be $t = t_1, ..., t_{12}$. Marginal Probabilities for a mutation in a branch is directly proportional to its numerator, i.e

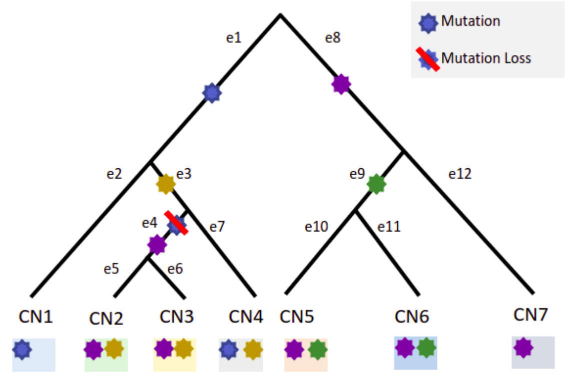


Fig. S2: Mutation Tree with all three type of mutations (Single, Back and Parallel) and leaves as clones

| Cell | $C_1$ | $C_2$ | $C_3$ | $C_4$ | $C_5$ | $C_6$ | $C_7$ | $C_8$ | $C_9$ | $C_{10}$ |
|---|---|---|---|---|---|---|---|---|---|---|
| Clone | $CN_1$ | $CN_1$ | $CN_2$ | $CN_3$ | $CN_4$ | $CN_5$ | $CN_5$ | $CN_6$ | $CN_6$ | $CN_7$ |

Fig. S3: Cell to Clone Mapping where each cell belong to a particular clone

| | $CN_1$ | | $CN_2$ | $CN_3$ | $CN_4$ | $CN_5$ | | $CN_6$ | | $CN_7$ |
|---|---|---|---|---|---|---|---|---|---|---|
| | $C_1$ | $C_2$ | $C_3$ | $C_4$ | $C_5$ | $C_6$ | $C_7$ | $C_8$ | $C_9$ | $C_{10}$ |
| $G_1$ | 1 | 1 | 0 | 0 | 1 | 0 | 0 | 0 | 0 | 0 |
| $G_2$ | 0 | 0 | 1 | 1 | 1 | 0 | 0 | 0 | 0 | 0 |
| $G_3$ | 0 | 0 | 1 | 1 | 0 | 1 | 1 | 1 | 1 | 1 |
| $G_4$ | 0 | 0 | 0 | 0 | 0 | 1 | 1 | 1 | 1 | 0 |

Fig. S4: True Genotype Matrix

| | $CN_1$ | | $CN_2$ | $CN_3$ | $CN_4$ | $CN_5$ | | $CN_6$ | | $CN_7$ |
|---|---|---|---|---|---|---|---|---|---|---|
| | $C_1$ | $C_2$ | $C_3$ | $C_4$ | $C_5$ | $C_6$ | $C_7$ | $C_8$ | $C_9$ | $C_{10}$ |
| $S_1$ | 1 | 1 | 0 | 0 | 1 | 0 | 0 | 1 | 0 | 0 |
| $S_2$ | 0 | 0 | 1 | 1 | 1 | 1 | 0 | 0 | 0 | 0 |
| $S_3$ | 0 | 1 | 1 | 0 | 0 | 1 | 0 | 1 | 1 | 1 |
| $S_4$ | 0 | 0 | 0 | 0 | 1 | 1 | 0 | 1 | 0 | 0 |

Fig. S5: Observed Genotype Matrix with Red showing inconsistency with True Genotype Matrix in fig. S4

1. Single Mutation (Green in $e_9$): The marginal probability that mutation $i$ occur in branch $e_9$ is

$$P(B_i^{0\to 1} = e_9 \mid \mathcal{T}, Q) \propto P_{00}(t_1)P_{00}(t_2)P_{00}(t_3)P_{00}(t_4)P_{00}(t_5)P_{00}(t_6)P_{00}(t_7)$$
$$P_{00}(t_8)P_{00}(t_{12})P_{10}(t_9)[P_{11}(t_{10})P_{11}(t_{11})]$$

2. Back Mutation (Blue in $e_1$ and $e_4$): The marginal probability that mutation $i$ occur

in branch $e_1$ and back mutate in branch $e_4$ is

$$P(B_i^{0\to1} = e_1, B_i^{1\to0} = e_4 \mid \mathcal{T}, Q) \propto P_{01}(t_1)[P_{11}(t_2)P_{11}(t_3)P_{11}(t_7)]P_{10}(t_4)[P_{00}(t_5)\\ P_{00}(t_6)][P_{00}(t_8)P_{00}(t_9)P_{00}(t_{10})P_{00}(t_{11})P_{00}(t_{12})]$$

Here $e_1$ and its descended $(e_2, e_3, e_4, e_5, e_6, e_7)$ are mutated with mutation $i$ and after back mutation, $e_4$ and its descended $(e_5, e_6)$ lost the mutation.

3. Parallel Mutation (Purple in $e_4$ and $e_8$): The marginal probability that mutation $i$ occur in branch $e_4$ and $e_8$ is

$$P(B_i^{0\to1} = e_1, B_i^{0\to1} = e_4 \mid \mathcal{T}, Q) \propto P_{00}(t_1)P_{00}(t_2)P_{00}(t_3)P_{00}(t_7)P_{01}(t_4)[P_{11}(t_5)\\ P_{11}(t_6)]P_{01}(t_8)[P_{11}(t_9)P_{11}(t_{10})P_{11}(t_{11})P_{11}(t_{12})]$$

Here $e_4$ and its descended $(e_5, e_6)$ are mutated with mutation i and also $e_8$ and its descended $(e_9, e_{10}, e_{11}, e_{12})$ have the mutation $i$.

#### 1.3 Marginal probability calculation example for Dollo model

An example of the marginal probability calculation for single and back mutations under Dollo model is given below.

For an example, fig. S8 and fig. S9 showing the observed and true binary genotype for mutations shown in tree (fig. S6). The set of branches is $E = e_1, ..., e_{16}$ and the corresponding set of branch lengths would be $t = t_1, ..., t_{16}$. Marginal Probabilities for a mutation in a branch is directly proportional to its numerator, i.e

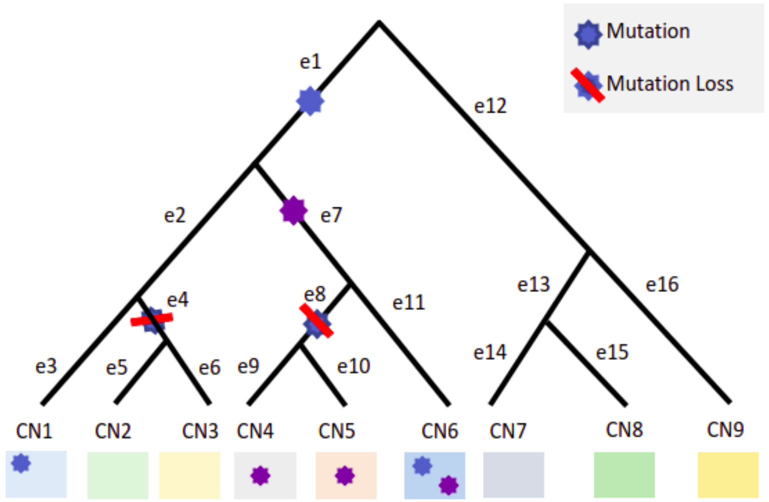


Fig. S6: Mutation Tree of Dollo Model with mutations (Single, Back) and leaves as clones

| Cell | $C_1$ | $C_2$ | $C_3$ | $C_4$ | $C_5$ | $C_6$ | $C_7$ | $C_8$ | $C_9$ | $C_{10}$ | $C_{11}$ | $C_{12}$ | $C_{13}$ |
|---|---|---|---|---|---|---|---|---|---|---|---|---|---|
| Clone | $CN_1$ | $CN_1$ | $CN_2$ | $CN_3$ | $CN_4$ | $CN_5$ | $CN_5$ | $CN_6$ | $CN_6$ | $CN_7$ | $CN_8$ | $CN_9$ | $CN_{10}$ |

Fig. S7: Cell to Clone Mapping where each cell belong to a particular clone

| | CN$_1$ | | CN$_2$ | CN$_3$ | CN$_4$ | CN$_5$ | | CN$_6$ | | CN$_7$ | CN$_8$ | CN$_9$ | CN$_{10}$ |
|---|---|---|---|---|---|---|---|---|---|---|---|---|---|
| | C$_1$ | C$_2$ | C$_3$ | C$_4$ | C$_5$ | C$_6$ | C$_7$ | C$_8$ | C$_9$ | C$_{10}$ | C$_{11}$ | C$_{12}$ | C$_{13}$ |
| G$_1$ | 1 | 1 | 0 | 0 | 0 | 0 | 0 | 1 | 1 | 0 | 0 | 0 | 0 |
| G$_2$ | 0 | 0 | 0 | 0 | 1 | 1 | 1 | 1 | 1 | 0 | 0 | 0 | 0 |

Fig. S8: True Genotype Matrix

| | CN$_1$ | | CN$_2$ | CN$_3$ | CN$_4$ | CN$_5$ | | CN$_6$ | | CN$_7$ | CN$_8$ | CN$_9$ | CN$_{10}$ |
|---|---|---|---|---|---|---|---|---|---|---|---|---|---|
| | C$_1$ | C$_2$ | C$_3$ | C$_4$ | C$_5$ | C$_6$ | C$_7$ | C$_8$ | C$_9$ | C$_{10}$ | C$_{11}$ | C$_{12}$ | C$_{13}$ |
| S$_1$ | 1 | 1 | 0 | 0 | 1 | 0 | 0 | 1 | 0 | 0 | 0 | 0 | 0 |
| S$_2$ | 0 | 1 | 0 | 0 | 1 | 1 | 0 | 1 | 1 | 0 | 0 | 0 | 1 |

Fig. S9: Observed Genotype Matrix with Red showing inconsistency with True Genotype Matrix in fig. S8

1. Single Mutation (Purple in $e_7$): The marginal probability that mutation $i$ occur in branch $e_9$ is

$$P(B_i^{0\to1} = e_7 \mid \mathcal{T}, Q) \propto P_{00}(t_1)P_{00}(t_2)P_{00}(t_3)P_{00}(t_4)P_{00}(t_5)P_{00}(t_6)$$
$$P_{00}(t_{12})P_{00}(t_{13})P_{00}(t_{14})P_{00}(t_{15})P_{00}(t_{16})P_{01}(t_7)$$
$$[P_{11}(t_8)P_{11}(t_9)P_{11}(t_{10})P_{11}(t_{11})]$$

2. Back Mutation (Blue in $e_1$, $e_4$ and $e_8$): The marginal probability that mutation $i$ occur in branch $e_1$ and back mutate in branch $e_4$ and $e_8$ is

$$P(B_i^{0\to1} = e_1, \{B_i^{1\to0} = y \mid y \in \{e_4, e_8\}\} \mid \mathcal{T}, Q) \propto P_{01}(t_1)[P_{11}(t_2)P_{11}(t_3)P_{11}(t_7)$$
$$P_{11}(t_{11})][P_{10}(t_4)P_{10}(t_8)][P_{00}(t_5)$$
$$P_{00}(t_6)][P_{00}(t_9)P_{00}(t_{10})][P_{00}(t_{12})$$
$$P_{00}(t_{13})P_{00}(t_{14})P_{00}(t_{15})P_{00}(t_{16})]$$

Here $e_1$ and its descended ($e_2, ..., e_{11}$) are mutated with mutation $i$ and after back mutation in $e_4$ and $e_8$, then $e_4$ and $e_8$ and all their descended ($e_5, e_6, e_9, e_{10}$) lost the mutation.

#### 1.4 Example for quantification of SCS error

An example of quantification of SCS error is given below.

Using the example in fig. S4 and fig. S5 the error probabilities for the three type of mutations can be written as

1. Single Mutation (Green in $e_9$): The probability of error when the true mutation $i = 1$ transitions from $0 \to 1$ on branch $e_9$ is

$$P(O_1 \mid B_i^{0\to1} = e_9, \mathcal{T}, N^i) \propto N_{00}^{11}N_{00}^{12}N_{00}^{13}N_{00}^{14}N_{01}^{15}$$
$$N_{11}^{16}N_{10}^{17}N_{11}^{18}N_{10}^{19}N_{00}^{110}$$

2. Back Mutation (Blue in $e_1$ and $e_4$): The probability of when the true mutation $i = 1$ transitions from $0 \to 1$ on branch $e_1$ and back mutation $1 \to 0$ on branch $e_4$ is

$$P(O_1 \mid B_i^{0\to1} = e_1, B_i^{0\to1} = e_4, \mathcal{T}, N^i) \propto N_{11}^{11}N_{11}^{12}N_{00}^{13}N_{00}^{14}N_{11}^{15}$$
$$N_{00}^{16}N_{00}^{17}N_{01}^{18}N_{00}^{19}N_{00}^{110}$$

3. Parallel Mutation (Purple in $e_4$ and $e_8$): The probability of error when the true mutation $i = 1$ transitions from $0 \to 1$ on branch $e_4$ and $e_8$ is

$$P(O_1 \mid B_i^{0\to1} = e_4, B_i^{0\to1} = e_8, \mathcal{T}, N^i) \propto N_{00}^{11} N_{01}^{12} N_{11}^{13} N_{10}^{14} N_{00}^{15} N_{11}^{16} N_{10}^{17} N_{11}^{18} N_{11}^{19} N_{11}^{110}$$

#### 1.5 Simulation Study

Settings for which we perform simulation study are given in Table S1. Setting 8 and 9 was used to simulated dataset under Infinite-site Assumption (ISA) and others datasets are simulated under Finite-site Assumption (FSA).

| Sr No | Clones | Cells | Sites | Clonal Mutation Rate | Recurrent Mutation Rate | FP | FN | d |
|---|---|---|---|---|---|---|---|---|
| 1 | 15 | 100 | 50 | 0.2 | 0.2 | 0.05 | 0.2 | NA |
| 2 | 15 | 100 | 50 | 0.2 | 0.2 | 0.05 | 0.3 | NA |
| 3 | 15 | 500 | 50 | 0.2 | 0.2 | 0.05 | 0.2 | NA |
| 4 | 15 | 500 | 50 | 0.2 | 0.2 | 0.05 | 0.3 | NA |
| 5 | 15 | 100 | 50 | 0.2 | 0.2 | 0.05 | 0.2 | 2 |
| 6 | 15 | 100 | 50 | 0.2 | 0.2 | 0.05 | 0.2 | 3 |
| 7 | 10 | 500 | 50 | 0.2 | 0.2 | 0.05 | 0.2 | NA |
| 8 | 15 | 100 | 50 | 0.2 | 0.2 | 0.05 | 0.2 | NA |
| 9 | 15 | 500 | 50 | 0.2 | 0.2 | 0.05 | 0.2 | NA |

Table S1: Parameter for Simulation Data.

##### 1.5.1 Simulation Process

To generate Simulation data we followed below steps

1. Generate 5 random bifurcating tumor clonal trees with $k$ clones by the recursive random splitting algorithm.
2. Randomly assign cells to clones of tree
3. Assign clonal mutation to first branch and remaining mutations are distributed on different branches i.e how many mutations are on particular branch
4. Randomly pick some sites and introduce recurrent mutations on these sites i.e. parallel mutations and back mutation. Number of recurrent sites are decided by recurrent rate
5. Record prefect true genotype data
6. Introduce noise to the true genotype data with error rate $\alpha$ and $\beta$, then record observed genotype data
7. In case of data for Dollo Model, back mutations are only recurrent mutations i.e. no parallel mutation

Repeat all steps with different choices of number of cells, error probabilities ($\alpha$ and $\beta$) and Dollo parameter $d$.

##### 1.5.2 Accuracy Estimate

We calculate accuracy of the MAP estimates across 5 tree for each setting using four different metrics as given below

**Location Accuracy**: To measure how accurately FiMO predict branches of mutation.

$$\frac{\text{Total number of mutation correctly inferred}}{\text{Total number of mutations}}$$

When inferred branch matches the true mutation branch then this inference is correct.

**Order Accuracy**: This is to measure order accuracy of all mutation pairs and here mutation pair need not be adjacent.

$$\frac{\text{Total number of mutation pairs whose order is correctly inferred}}{\text{Total number of true mutation pairs}}$$

Correctly inferred mutation means, If one pair of mutations is acquired on two branches in a simulated tree, then the inferred mutation order on any two branches must be the same.

**Adjacent Order Accuracy**: This measure is similar to order accuracy but here we taken adjacent pairs only.

$$\frac{\text{Total number of adjacent mutation pairs whose order is correctly inferred}}{\text{Total number of true adjacent mutation pairs}}$$

Correctly inferred mutation means, If one pair of mutations is acquired on two adjacent branches in a simulated tree, then the inferred mutation order on any two adjacent branches must be the same.

**Partition Accuracy**: This Accuracy measure is useful when topology of true tree and observed tree are different. It works at cell level and give idea of state of how many cells are correctly predicted.

$$\frac{\text{Accuracy of mutated cells + Accuracy of non mutated cells}}{2}$$

where,

$$\text{Accuracy of mutated cells} = \frac{\text{Number of cell mutated in both true and observed tree}}{\text{Total number of cell mutated in either of tree}}$$

Accuracy of mutated cell gives idea of cells that are in mutated state and common to both true tree and observed tree.

$$\text{Accuracy of non mutated cells} = \frac{\text{Number of cell non mutated in both true and observed tree}}{\text{Total number of cell non mutated in either of tree}}$$

Accuracy of non mutated cell gives idea of cells that are in normal state and common to both true tree and observed tree.

### 2 Supplementary Results

Average accuracy for each setting is given in Table S2

| Setting | Location Accuracy | Order Accuracy | Adjacent Order Accuracy | Partition Accuracy |
|---|---|---|---|---|
| 1 | 94.0 | 93.9 | 84.5 | 98.6 |
| 2 | 88.8 | 90.9 | 85.5 | 96.3 |
| 3 | 99.6 | 99.9 | 99.8 | 99.9 |
| 4 | 96.0 | 95.5 | 90.6 | 99.5 |
| 5 | 89.6 | 85.9 | 76.0 | 97.9 |
| 6 | 85.2 | 81.6 | 73.2 | 96.6 |
| 7 | 99.2 | 98.7 | 98.4 | 99.9 |
| 8 | 98.8 | 99.9 | 98.6 | 99.7 |
| 9 | 99.2 | 99.8 | 98.4 | 99.8 |

Table S2: Average Accuracy of FiMO for all settings

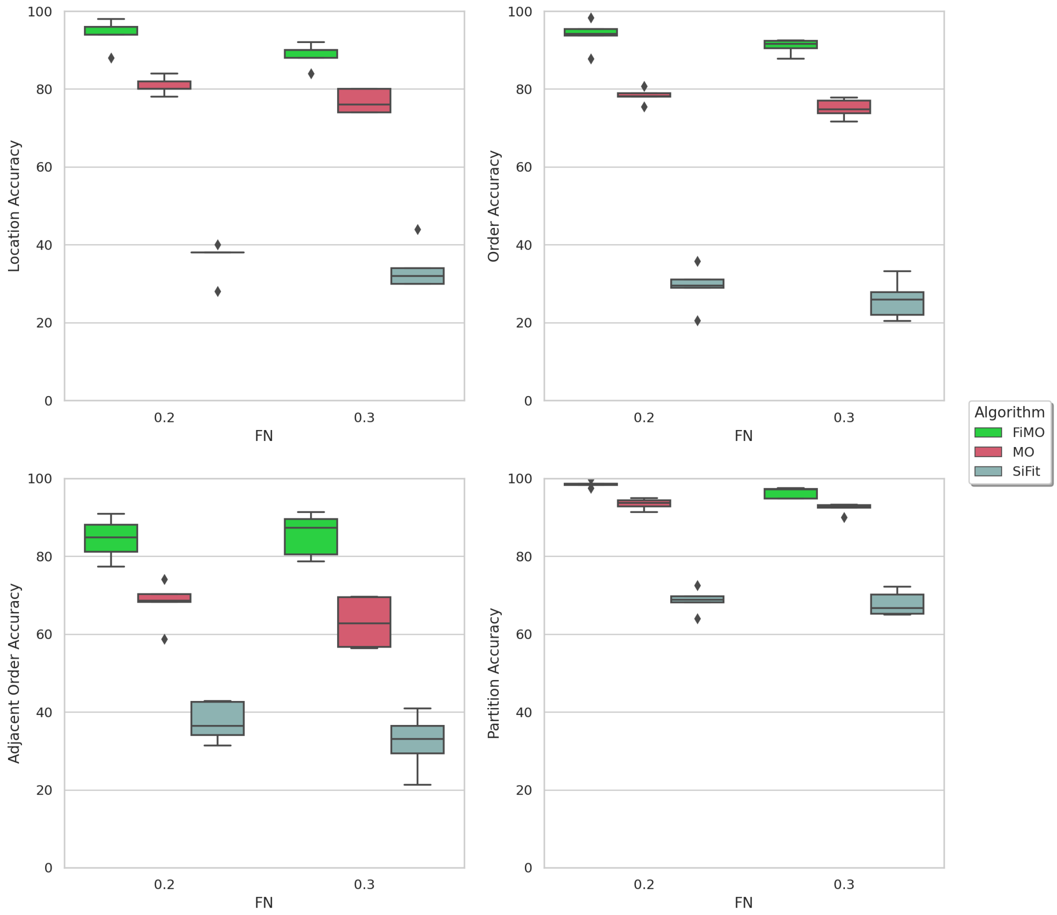


Fig. S10: Performance comparison of FiMO, MO and SiFit for varying false negative rate for 100 cells datasets. False negative rate was varied $\beta \in \{0.2, 0.3\}$. The y-axes show different accuracy measures.

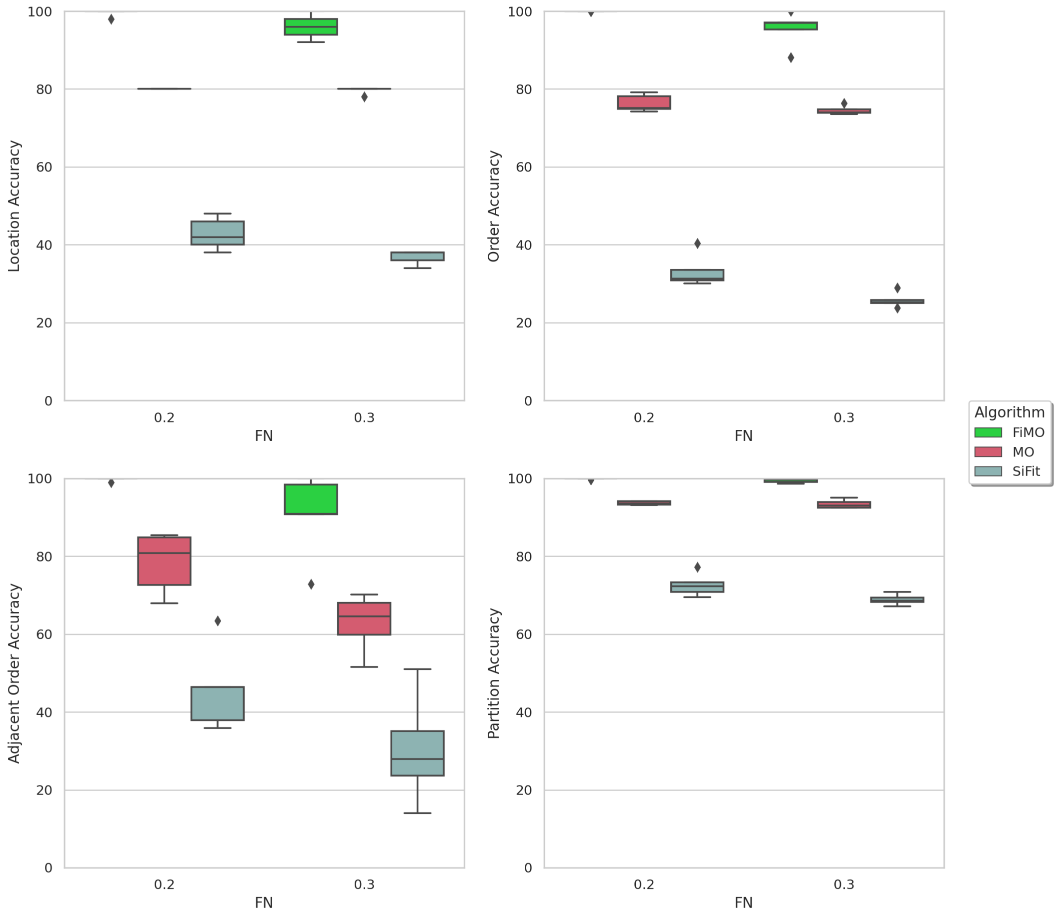


Fig. S11: Performance comparison of simulated datasets containing 500 cells. FiMO performance is compared against MO and SiFit on simulated datasets containing 500 cells for varying FN error rate. The y-axes show different accuracy measures.

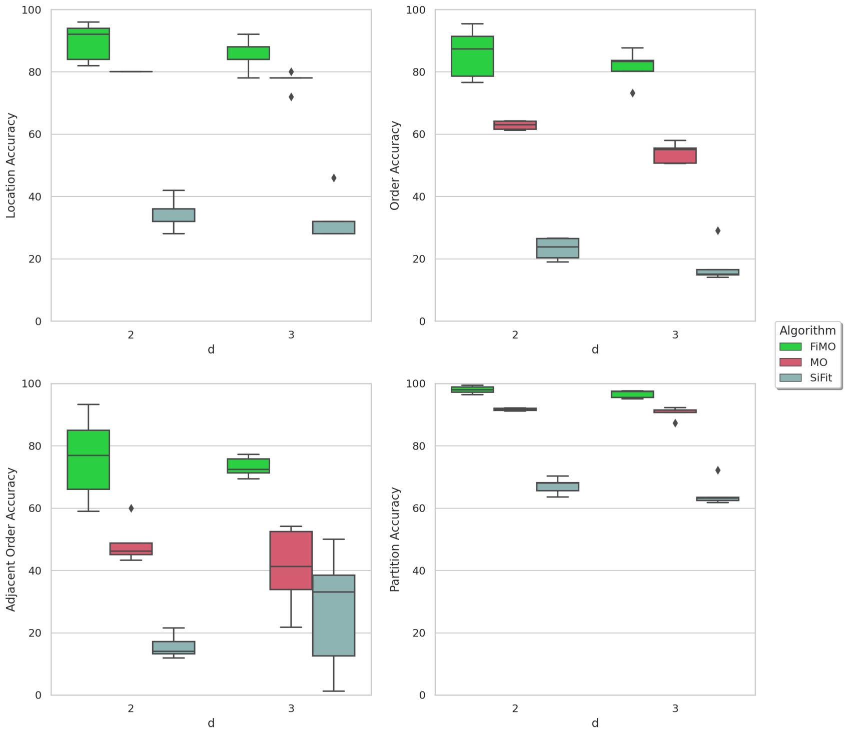


Fig. S12: Performance comparison of simulated datasets having 0.2 FN and 100 cells. FiMO performance is compared with MO and SiFit on simulated datasets having 0.2 FN, 100 cells and varying for d (Maximum number of back mutation for a site). On the x-axis, we have d and On the y-axis, Different Accuracy measures are plotted.

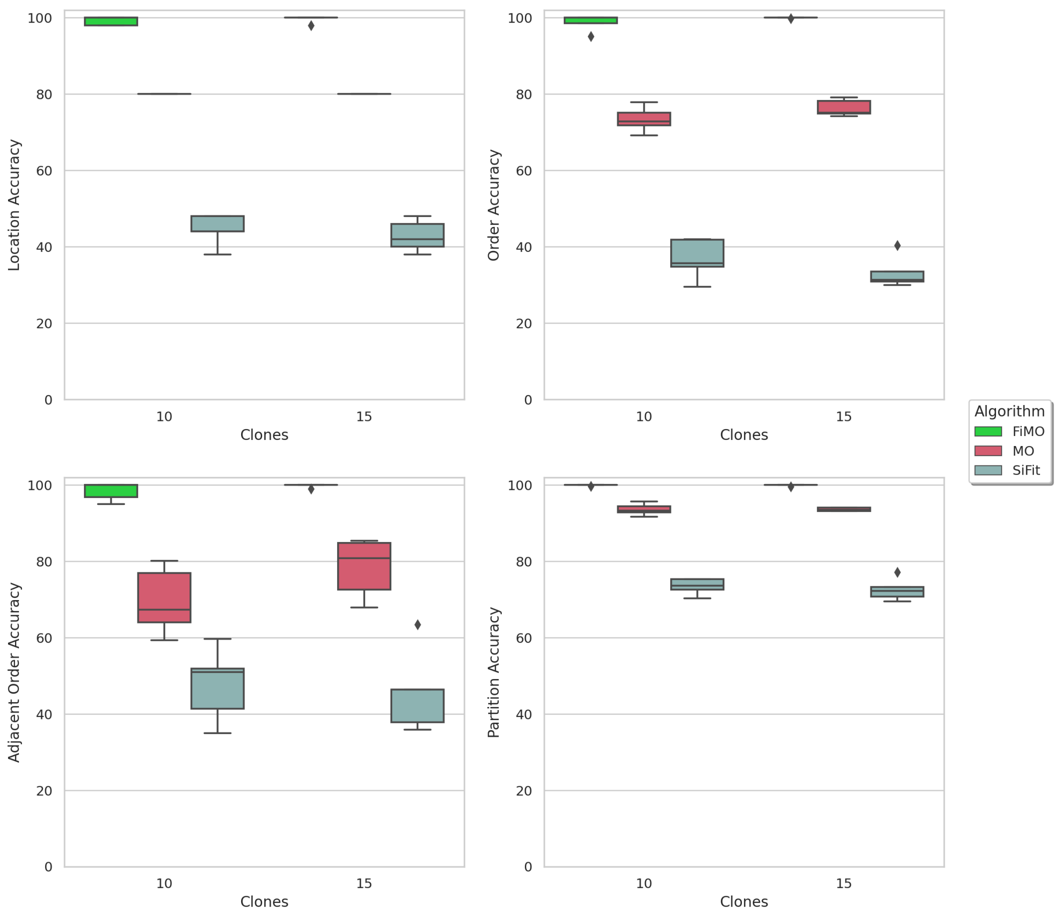


Fig. S13: Performance comparison of FiMO, MO and SiFit for varying number of clones. Clones was varied $k \in \{10, 15\}$. The y-axes show different accuracy measures.

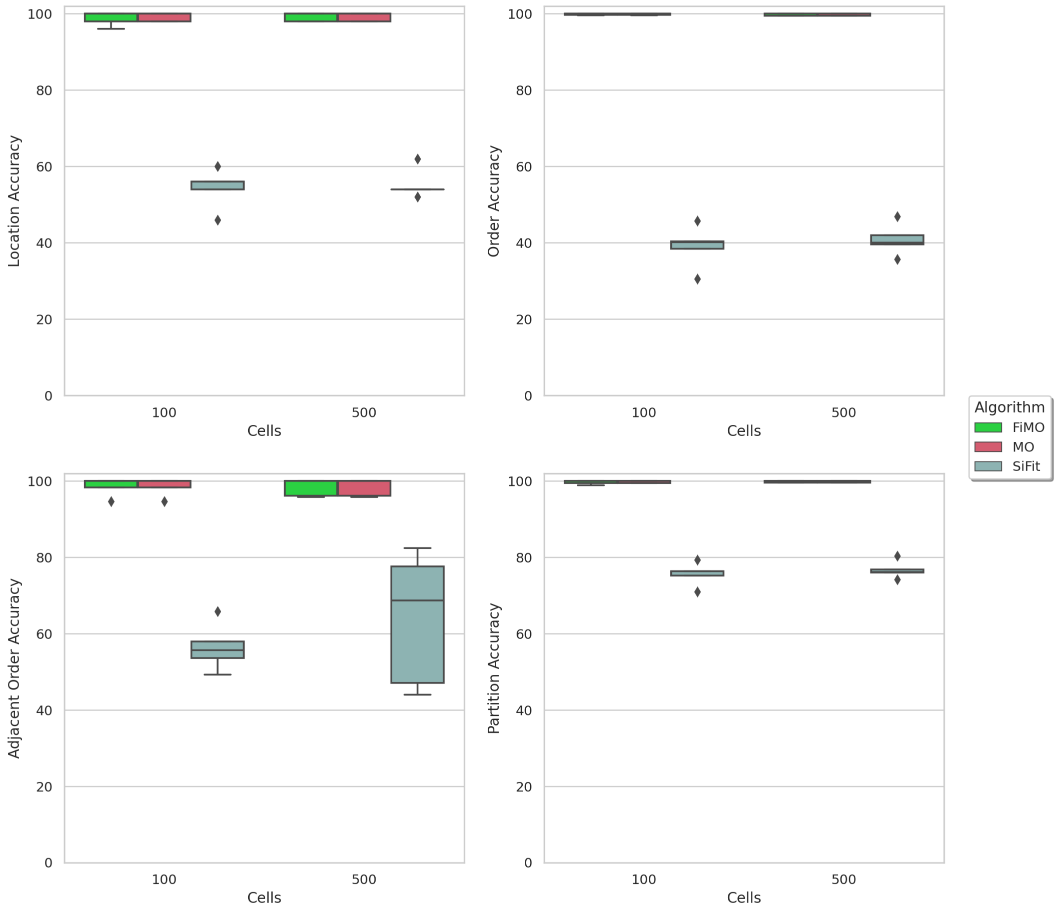


Fig. S14: Performance comparison of FiMO, MO and SiFit for varying number of cell. Cells was varied $m \in \{100, 500\}$ and dataset used for this comparison was simulated under infinite-site assumption. The y-axes show different accuracy measures.
